# Two Stages of Speech Envelope Tracking in Human Auditory Cortex Modulated by Speech Intelligibility

**DOI:** 10.1101/2021.12.11.472249

**Authors:** Na Xu, Baotian Zhao, Lu Luo, Kai Zhang, Xiaoqiu Shao, Guoming Luan, Qian Wang, Wenhan Hu, Qun Wang

**Affiliations:** Department of Neurology, Beijing Tiantan Hospital, Capital Medical University, Beijing 100070, China; Department of Neurosurgery, Beijing Tiantan Hospital, Capital Medical University, Beijing 100070, China; Beijing Neurosurgical Institute, Capital Medical University, Beijing 100070, China; School of Psychology, Beijing Sport University, Beijing 100084, China; Beijing Key Laboratory of Epilepsy, Epilepsy Center, Sanbo Brain Hospital, Capital Medical University, Beijing 100093, China; School of Psychological and Cognitive Sciences, Beijing Key Laboratory of Behavior and Mental Health, Peking University, Beijing 100871, China; National Clinical Research Center for Neurological Diseases, Beijing 100070, China; Beijing Institute of Brain Disorders, Collaborative Innovation Center for Brain Disorders, Capital Medical University, Beijing 100069, China; IDG/McGovern Institute for Brain Research, Peking University, Beijing 100871, China

**Keywords:** stereo-electroencephalogram (sEEG), envelope tracking, high-γ band power, θ band phase, spectrally degraded speech

## Abstract

The envelope is essential for speech perception. Recent studies have shown that cortical activity can track the acoustic envelope. However, whether the tracking strength reflects the extent of speech intelligibility processing remains controversial. Here, using stereo-electroencephalogram (sEEG) technology, we directly recorded the activity in human auditory cortex while subjects listened to either natural or noise-vocoded speech. These two stimuli have approximately identical envelopes, but the noise-vocoded speech does not have speech intelligibility. We found two stages of envelope tracking in auditory cortex: an early high-γ (60-140 Hz) power stage (delay ≈ 49 ms) that preferred the noise-vocoded speech, and a late θ (4-8 Hz) phase stage (delay ≈ 178 ms) that preferred the natural speech. Furthermore, the decoding performance of high-γ power was better in primary auditory cortex than in non-primary auditory cortex, consistent with its short tracking delay. We also found distinct lateralization effects: high-γ power envelope tracking dominated left auditory cortex, while θ phase showed better decoding performance in right auditory cortex. In sum, we suggested a functional dissociation between high-γ power and θ phase: the former reflects fast and automatic processing of brief acoustic features, while the latter correlates to slow build-up processing facilitated by speech intelligibility.

## INTRODUCTION

The temporal envelopes of sounds contain critical information for speech understanding (Shannon et al., 1995; Smith et al., 2002). When listening to speech, cortical activities, especially those in auditory cortex, can temporally track the acoustic envelopes (Poeppel and Assaneo, 2020). The strength of cortical envelope tracking may reflect the extent of speech processing (Di Liberto et al., 2018; Etard and Reichenbach, 2019; Vanthornhout et al., 2018). However, acoustic signals with no speech intelligibility (i.e., time-reversed speech or music) could also be well tracked by the cortical activities (Doelling and Poeppel, 2015; Harding et al., 2019; Howard and Poeppel, 2010). Therefore, whether cortical envelope tracking functionally contributed to speech intelligibility remains unclear.

To address this long-standing issue, an idea is to test the cortical activities to sounds with identical temporal envelopes but different intelligibility. Following this line of thought, a noise-vocoding technique can degrade the spectral regularity while preserving the speech envelope (Davis et al., 2005). Moreover, varying the number of frequency bands of the vocoder modulates the speech intelligibility effectively: with decreasing number of frequency bands, the intelligibility of the noise-vocoded speech was gradually destroyed (Davis and Johnsrude, 2003). However, how intelligibility level modulates cortical tracking of noise-vocoded speech remained puzzled. Some previous studies showed that cortical envelope tracking strength enhanced with increased speech intelligibility (Ding et al., 2014; Peelle et al., 2013), while others found not (Millman et al., 2015; Rimmele et al., 2015). In a recent study, Hauswald and collages (2020) found an inverted U-shaped relationship between the cortical envelope tracking strength and speech intelligibility.

These inconsistent findings implied the existence of different neural mechanisms of speech envelope tracking in auditory cortex, which has been rarely addressed. Electrophysiological evidence showed that both low and high-frequency cortical activities could track speech envelope (Gourevitch et al., 2020). In low-frequency bands, one of the most robust observations was that envelope could be tracked by the phase in θ band (4-8 Hz). On the other hand, the high-frequency envelope tracking was mainly contributed by the fluctuated power in high-γ band (~70-150 Hz) (Kubanek et al., 2013; Kulasingham et al., 2020; Nourski et al., 2009; Synigal et al., 2020; Zion Golumbic et al., 2013). Both θ phase envelope tracking and high-γ power ould be modulated by noise-vocoded speech in different intelligibility levels (Ding et al., 2014; Hauswald et al., 2020; Nourski et al., 2019; Peelle et al., 2013). However, there is still a lack of direct comparison between θ phase and high-γ power tracking patterns in auditory cortex, hindering our understanding of their roles in representing speech intelligibility information.

Using stereo-electroencephalogram (sEEG) technology in ten patients with refractory epilepsy, we directly recorded the electrophysiological signals in human auditory cortex when patients were passively listening to a natural or a two-band noise-vocoded speech. In the single-contact level, we analyzed the envelope tracking strength, tracking delay, and decoding accuracy of both high-γ (60-140 Hz) power and θ phase (4-8 Hz) in Heschl’s gyrus (HG) and superior temporal gyrus (STG). HG includes mostly the primary auditory cortex, and STG is considered the non-primary auditory cortex (Clarke and Morosan, 2012). We then compared the tracking pattern and decoding performance in different hemispheres and brain areas. Our results revealed distinct processing stages of speech intelligibility subserved by high-γ power and θ phase in auditory cortex.

## RESULTS

### Auditory responsive contacts

We first selected the sEEG contacts in Heschl’s gyrus (HG) and superior temporal gyrus (STG) in all ten patients. Out of 115 contacts, 75 contacts (HG, n = 20; STG, n = 55) (Figure 2A) with both significant auditory evoked high-γ signal and broadband signal were further analyzed.

**Figure 1.**
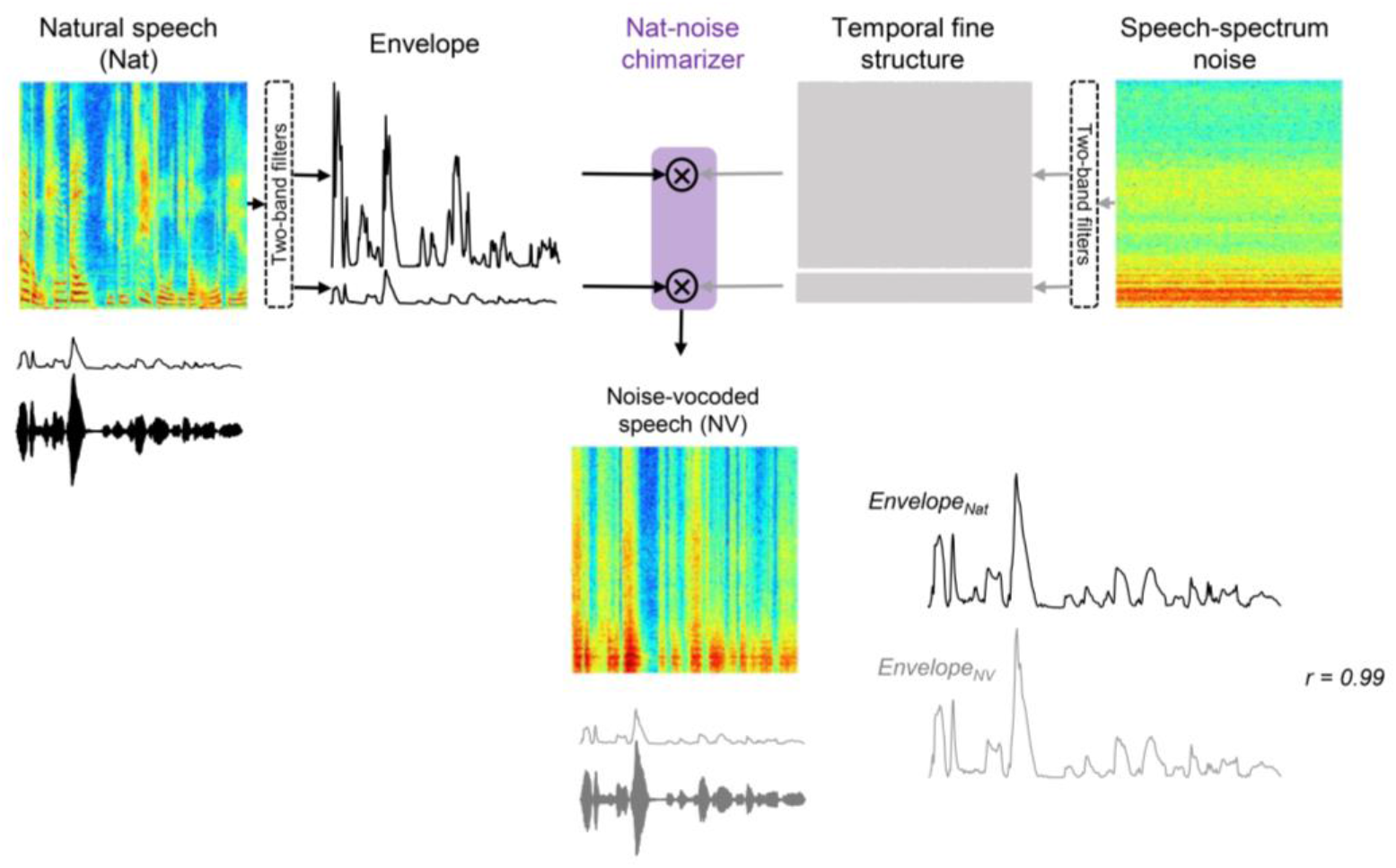
Schematic illustration of the noise-vocoded speech synthesis. Firstly, the natural speech and the speech-spectrum noise are filtered into two bands. Secondly, the outputs of these two banks are Hilbert transformed to extract the envelopes and the temporal fine structures (TFSs). Thirdly, the envelope of the natural speech is multiplied by the TFS of the speech-spectrum noise in each band. Finally, these two chimeras are summed to form the noise-vocoded speech (NV). Notably, the envelopes of the natural speech and that of the noise-vocoded speech are approximately identical (correlation coefficient = 0.99). *Nat, natural speech; NV, noise-vocoded speech*.

**Figure 2.**
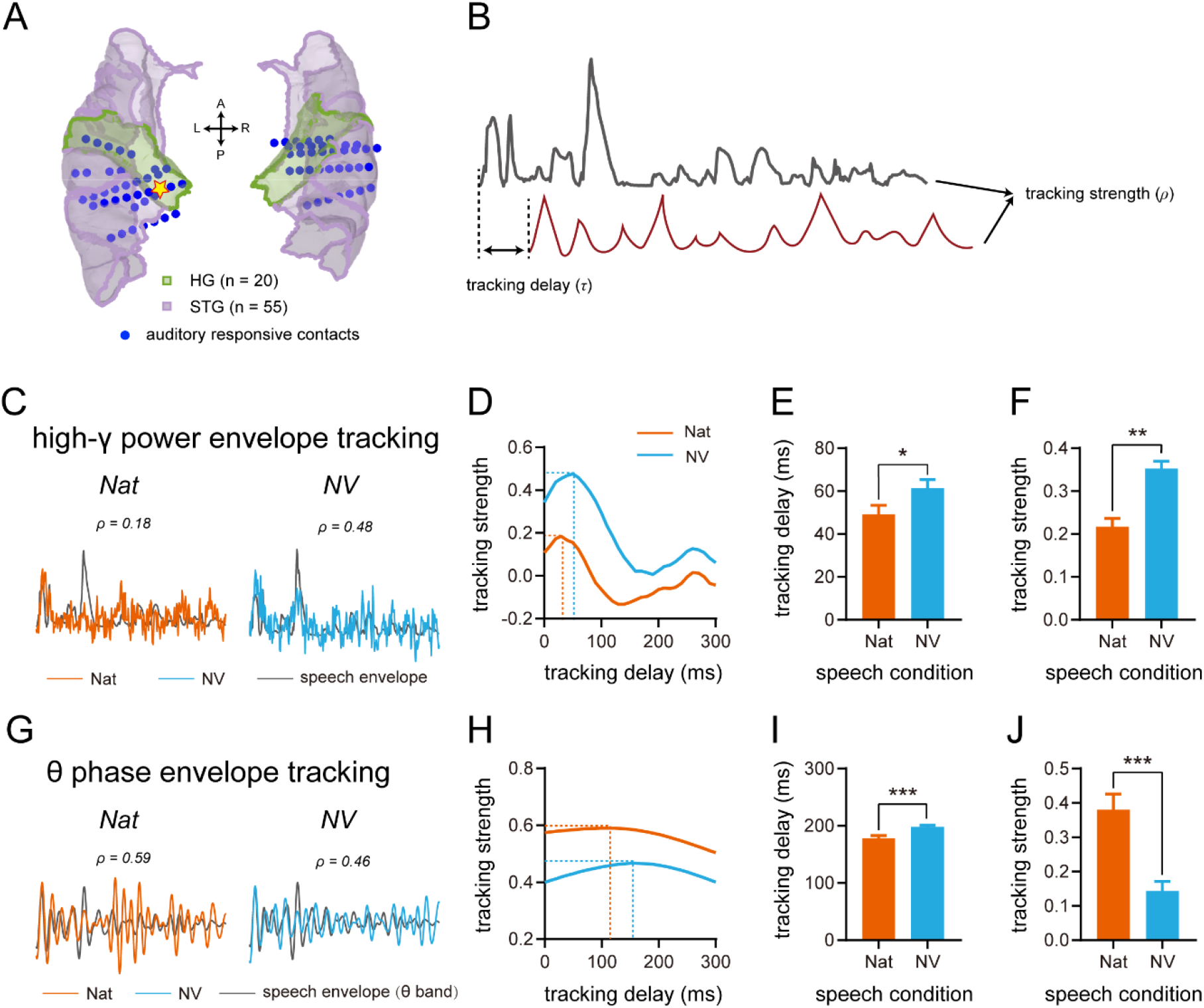
**A**, anatomical parcellation of temporal lobe regions of the human auditory cortex, and contacts selected from all patients counts across the selected anatomical areas and projected onto a Montreal Neurological Institute (MNI) atlas brain (BCI-DNI_BrainSuite_2016). HG, Heschl’s gyrus (in green); STG, superior temporal gyrus (in purple), the exemplary contact is marked by a yellow five-pointed star with the red edge. **B**, the diagram showing the tracking strength (*ρ*) of two example waveforms obtained at the optimal tracking delay (τ). **C**, the waveform of the Nat-speech envelope and the NV-speech envelope (gray curves), and the corresponding averaged high-γ power responses (Nat, orange curves; NV, blue curves) of the example contact (five-pointed star in A). **D**, high-γ power envelope tracking for Nat-speech and NV-speech at different lag times are plotted for the example waveforms shown in **C**. The dashed lines show the maximum tracking strengths at the optimal delay times. **E**, comparison of the tracking delay for high-γ power between the condition of Nat-speech and NV-speech. **F**, comparison of tracking strength for high-γ power between Nat-speech and NV-speech. G, single-trial θ band responses of the example contact (five-pointed star in A) for Nat-speech (orange curves) and NV-speech (blue curves), and the waveforms of the speech envelope (gray curves) filtered by θ band (4-8 Hz). **H**, θ phase envelope tracking for Nat-speech and NV-speech at different lag times are plotted for the example waveforms shown in **G**. **I**, tracking delay of θ phase for Nat-speech and NV-speech is compared. **J**, θ phase tracking strength is compared for Nat-speech and NV-speech. *Nat, natural speech; NV, noise-vocoded speech*.

### Early high-γ power envelope tracking preferred the noise-vocoded speech

The patients passively listened to a natural speech and a noise-vocoded speech (duration = 3.275 s), each with a repetition of 30 times. In each sEEG contact, we analyzed the envelope tracking strength (ρ) and delay (τ) by conducting a cross-correlation test between time series of neural high-γ power and acoustic envelope (Figure 2B).

Figure 2C showed waveforms of the envelopes and high-γ power under natural and noise-vocoded speech conditions from one representative contact. As shown in Figure 2D, we estimated the optimal tracking strength and tracking delay under each speech condition at the single-contact level, where a positive lag time indicates that high-γ power lags the speech envelope. The envelope tracking strength peaked with a 30 ms optimal tracking delay for natural speech and a 50 ms optimal tracking delay for noise-vocoded speech (natural ρ = 0.18; noise vocoded ρ = 0.48; Figure 2D). We tested the statistical significance of high-γ power envelope tracking strength for each contact. Seventy-two contacts (out of 75 contacts) showed significant tracking and were kept for further comparisons. We compared both the optimal tracking delay and strength of high-γ power under two speech conditions using paired *t*-tests. The tracking delay for natural speech was significantly shorter than that for noise-vocoded speech (natural: 49.2 ± 4.3 ms, mean ± SE; noise-vocoded: 61.4 ± 4.0 ms; *t*_71_ = −2.146, *p* = 0.035), while the tracking strength for natural speech condition was significantly lower than that for noise-vocoded speech (natural: 0.22 ± 0.02; noise-vocoded: 0.35 ± 0.02; *t*_71_ = 10.909, *p* < 0.001; Figure 2E and F). These results revealed that the high-γ power tracked the speech envelope in the early stage of auditory processing (< 100 ms), and the tracking strength was enhanced when speech intelligibility was destroyed.

### Late θ phase envelope tracking preferred the natural speech

We also analyzed the envelope tracking strength (ρ) and delay (τ) between neural θ phase and θ band-filtered envelope phase along the time series using the cerebro-acoustic coherence (CACoh) index (Harding et al., 2019; Peelle et al., 2013). Figure 2G showed waveforms of the θ band-filtered envelope and the neural θ signal under natural and noise-vocoded speech conditions, extracted from the same contact in Figure 2C. Similar to high-γ power tracking, we estimated the peak tracking strength and optimal tracking delay of θ phase under each speech condition at the single-contact level, where a positive lag time indicates that θ phase lags the speech envelope (Figure 2H). For each contact, we built a null distribution for θ phase envelope tracking by permutation (100 times) to examine its significance. Forty-six contacts (out of 75 contacts) were significant and included in the further analysis. We compared the optimal tracking delay and strength of the θ phase under two speech conditions using paired *t*-tests. We found that the θ phase tracking delay for natural speech was significantly shorter than that for noise-vocoded speech (natural: 178.1 ± 5.0 ms; noise-vocoded: 198.2 ±3.1 ms; *t*_45_ = −3.903,*p* < 0.001), while the tracking strength for natural speech was significantly larger than that for noise-vocoded speech (natural: 0.38 ± 0.05; noise-vocoded: 0.14 ± 0.03; *t*_45_ = 5.882, *p* < 0.001; Figure 2I and J). These results revealed that the θ phase tracked the speech envelope in the late stage of auditory processing (> 100 ms), and the tracking strength decreased when speech intelligibility was destroyed.

### Left hemisphere lateralization of high-γ power envelope tracking

Previous findings suggest that high-frequency and low-frequency neural oscillations in the human auditory cortex showed functional lateralization in hemispheres (Giroud et al., 2020; Morillon et al., 2012; Poeppel, 2003) and functional dominance in brain areas (Fontolan et al., 2014). Thus, we investigated whether the envelope tracking strength of high-γ power and θ phase also showed lateralization or brain area effects. We pooled the data across natural and noise-vocoded speech conditions, and test the difference between hemisphere/brain areas using *t*-tests. Contacts with strong high-γ power tracking strength clustered in the left hemisphere (Figure 3A), and *t*-tests results confirmed that the envelope tracking strength of high-γ power of the left hemisphere was significantly larger than that of the right hemisphere (left: 0.36 ± 0.02; right: 0.20 ± 0.01; *t*_142_ = 6.450, *p* < 0.001) (Figure 3B). However, we did not observe significant difference for high-γ power envelope tracking between brain areas (HG: 0.27 ± 0.03; STG: 0.28 ± 0.02; *t*_142_ = −0.550, *p* = 0.583) (Figure 3B). These results suggested a left hemisphere lateralization for high-γ power envelope tracking.

**Figure 3.**
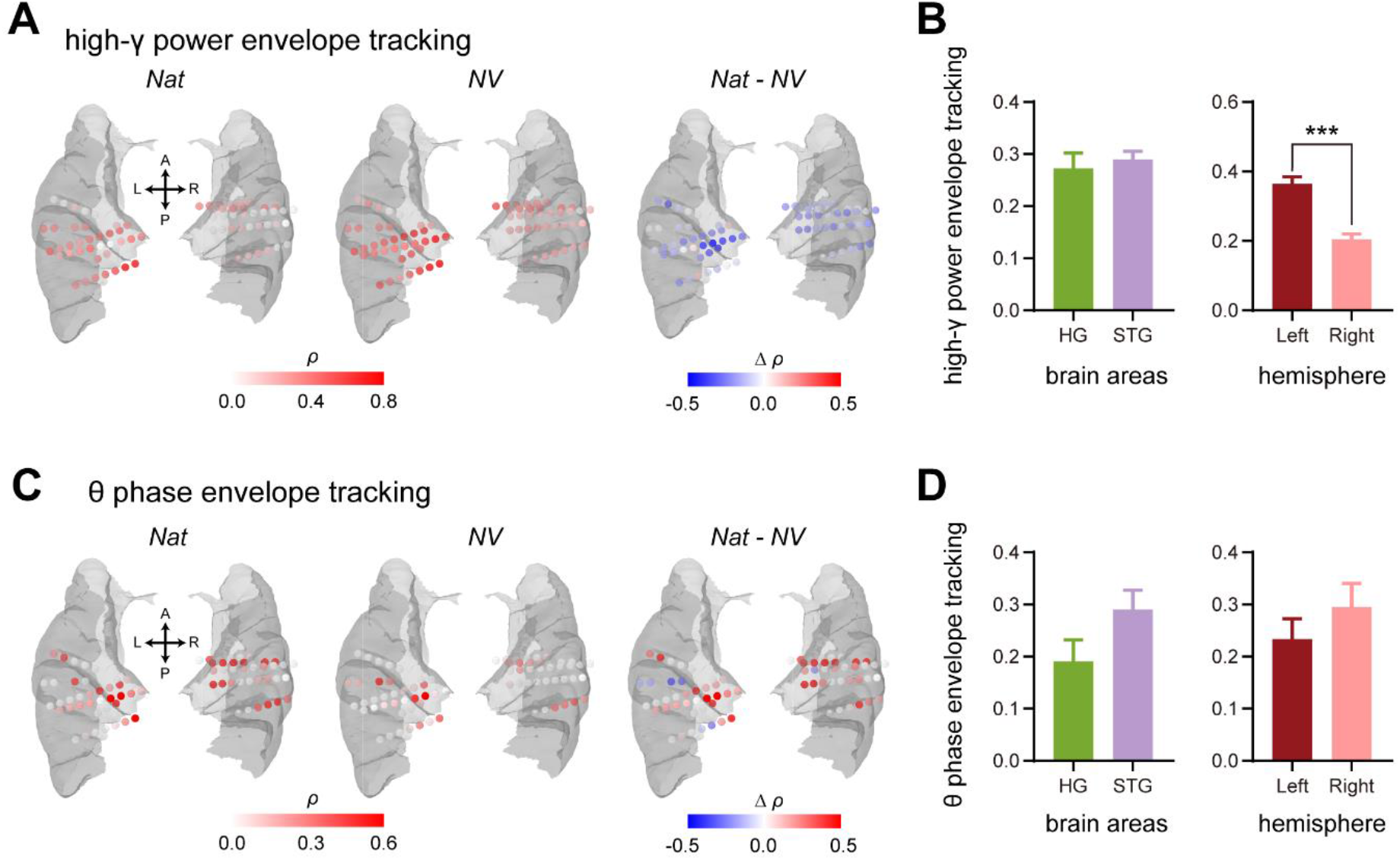
Envelope tracking in high-γ power and θ phase. **A**, high-γ power envelope tracking for Nat-speech (left subpanel), NV-speech (middle subpanel), and the difference between these two conditions (Nat-NV) (right subpanel) are plotted on the brain areas selected. **B**, Comparison of the high-γ power envelope tracking for the speech stimuli (Nat- and NV-) between brain areas (left subpanel, HG vs. STG), and between left and right hemispheres (right subpanel). **C**, θ phase envelope tracking for Nat-speech (left subpanel), NV-speech (middle subpanel), and the difference between these two conditions (Nat-NV) (right subpanel) are plotted on the brain. **D**, θ phase envelope tracking is compared for the speech stimuli (Nat- and NV-) between brain areas (left subpanel, HG vs. STG) and between left and right hemispheres (right subpanel). *Nat, natural speech; NV, noise-vocoded speech*. \*\*\**p* < 0.001.

Figure 3C showed the θ phase envelope tracking strength of single contacts. θ phase envelope tracking between was also compared between hemisphere/brain areas, and the results of *t*-tests showed no significant difference in HG and STG (HG: 0.19 ± 0.04; STG: 0.29 ± 0.04; *t*_90_ = −1.523, *p* = 0.131), and no significant difference between left and right hemispheres (left: 0.23 ± 0.04; right: 0.30 ± 0.04; *t*_90_ = −1.039,*p* = 0.301) (Figure 3D).

### Distinct decoding performance of high-γ power and the θ phase

By far, our results suggest that high-γ power and θ phase tracked the envelopes of natural and noise-vocoded speech with different tracking strengths and delays. To examine whether these tracking patterns reflected the acoustic information representation in auditory cortex, we decoded the speech condition with single-trial high-γ power or the θ phase for each contact using the support vector machine (SVM) approach via leave-one-out cross-validations.

For high-γ power, speech condition could be decoded in 49% of the contacts (statistical significance above the chance level). Chi-square tests showed that the HG had higher proportions of significant decoded contacts than STG (HG: 80.0%; STG: 38.2%; χ^2^= 10.261, *p* = 0.001), while no significance was found between left and right hemisphere (left: 50.0%; right: 48.6%; χ^2^ = 0.014, *p* = 0.907; Figure 4B). We further testes the decoding accuracy between different brain areas and different hemispheres. The results revealed that high-γ power decoding accuracy in HG contacts were significantly higher than that in STG contacts (HG: 0.76 ± 0.03; STG: 0.63 ± 0.02; *t*_73_ = 3.270, *p* = 0.002), but no significance was found between left and right hemisphere (left: 0.68 ± 0.02; right: 0.65 ± 0.03; *t*_73_ = 0.657, *p* = 0.513; Figure 4C). These results indicated that high-γ power decoding performance was predominant in Heschls’ gyrus.

**Figure 4.**
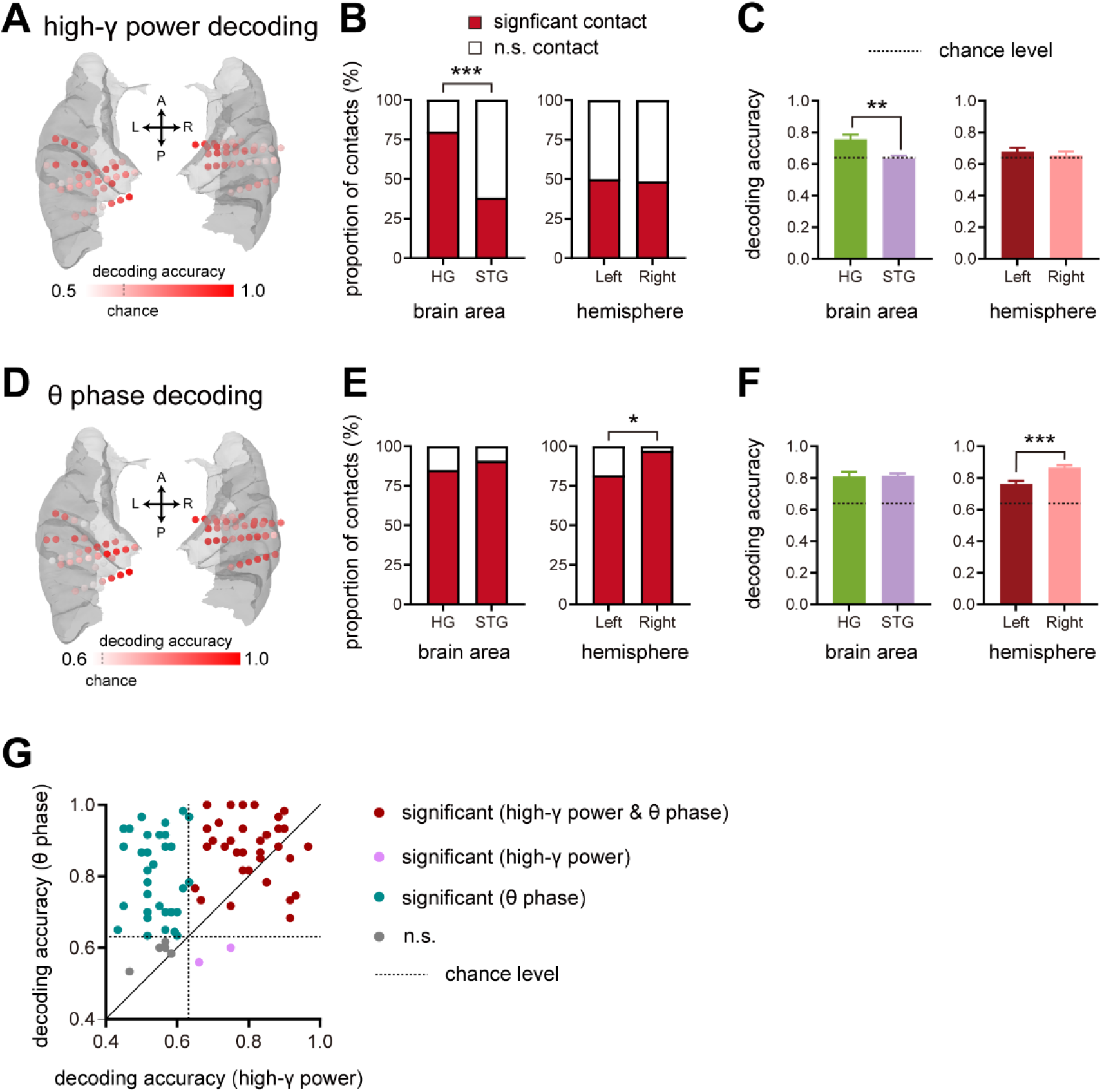
Decoding of Nat-speech and NV-speech using the feature of either high-γ power or θ phase. **A,** distributions of high-γ power decoding accuracy for each contact in the selected brain areas. **B**, proportions of the significant and the non-significant (n.s.) contacts for high-γ power decoding accuracy across brain areas and hemispheres, respectively. **C**, comparisons of high-γ power decoding accuracy between brain areas (left subpanel, HG vs. STG) and between left and right hemispheres (right subpanel), respectively. **D,** distributions of θ phase decoding accuracy for each contact in the selected brain areas. **E**, proportions of the significant and the non-significant (n.s.) contacts for θ phase decoding accuracy across brain areas and hemispheres, respectively. **F**, θ phase decoding accuracy between brain areas (left subpanel, HG vs. STG) and between left and right hemispheres (right subpanel) are compared respectively. **G,** scatter plot shows decoding accuracy using high-γ power feature against the decoding accuracy using θ phase feature for each contact. Values with different colors labeled show the statistical significance of their decoding accuracy. Dotted lines in this figure denote the chance level obtained by permutation testing (n = 1000). Decoding values above these dotted lines are higher than 95% of random values. *\*\*\*p* < 0.001, *\*\*p* < 0.01, *\*p* < 0.05.

For θ phase, speech condition could be decoded in 89% of the contacts. Chi-square tests showed that the right hemisphere had higher proportions of significant decoded contacts than the left hemisphere (left: 81.6%; right: 97.3%; χ^2^ = 4.861, *p* = 0.027), while no significance was found between HG and STG (HG: 85.0%; STG: 90.9%; χ^2^ = 0.537, *p* = 0.463; Figure 4E). We then testes the decoding accuracy between different brain areas and different hemispheres. The results revealed that θ phase decoding accuracy in right hemisphere were significantly higher than that in left hemisphere (left: 0.76 ± 0.02; right: 0.86 ± 0.02; *t*_73_ = −3.842, *p* < 0.001), but no significance was found between HG and STG (HG: 0.81 ± 0.03; STG: 0.81 ± 0.02; *t*_73_ = −0.104, *p* = 0.917; Figure 4F). These results suggested a right hemisphere dominance of θ phase decoding performance.

Finally, we compared the decoding performance of high-γ power versus θ phase and found that not only the proportion of significant contacts was larger when using θ phase (χ^2^ = 28.219, *p* < 0.001), but also the mean decoding accuracy using θ phase was better than that using high-γ power (θ phase: 0.81 ± 0.01; high-γ power: 0.67 ± 0.02; *t*_74_ = 7.706, *p* < 0.001; Figure 4G).

## DISCUSSION

Using sEEG recordings, we revealed two distinct processing stages of speech envelope in human auditory cortex modulated by speech intelligibility. In the early stage, high-γ power tracked speech envelope with a short latency (≈ 49 ms) and enhanced when the speech sound was incomprehensible. In the late stage, θ phase tracked speech envelope with a long latency (≈ 178 ms) and decreased with degrading speech intelligibility. Consistent with the tracking latency, we also found that the spectra-temporal information in high-γ power represented better in primary auditory cortex than in non-primary auditory cortex. Furthermore, we observed the different hemisphere lateralization of high-γ power and θ phase, suggesting that these two neural activities may have highly specific linguistic significance.

### Early high-γ power and late θ phase envelope tracking

We showed reliable envelope tracking in high-γ power and θ phase differed in their tracking delay. The relative short tracking delay (about 50 ms) of high-γ power is in line with prior work (Nourski et al., 2014; Zion Golumbic et al., 2013). In contrast, θ phase tracking delay is long (>100 ms), which is also consistent with previous studies (Di Liberto et al., 2015; Harding et al., 2019; Zou et al., 2021). And the finding that both of high-γ power and θ phase showed the shorter tracking delay for the natural speech compared with that for the noise-vocoded speech, suggests the sensitivity to natural human speech (Yu and Wang, 2021). Sound encoding is a hierarchical, feedforward processing framework in human auditory cortex (Hickok and Poeppel, 2007). The distinct tracking delays of high-γ power and θ phase indicate they may be involved in different stages of speech processing. The tracking delay in high-γ power is similar to the evoked onset response of human auditory cortex (Elhilali et al., 2004). Moreover, we also revealed a better decoding performance using high-γ power feature clustered in Heshl’s gyrus, which suggests the high-γ power envelope tracking is an early response in primary auditory cortex. We did not observed a significant effect for brain areas for θ phase decoding performance (Figure 4). However, we found a trend of larger θ phase tracking strength in STG than Heshl’s gyrus (Figure 3).

### Distinct functional roles of high-γ power and θ phase

It has been suggested that there may be different functional roles for neural activities at different frequencies in speech processing (Mai et al., 2016). A previous ECoG study also showed a distinction between high-γ power and low-frequency in tracking the envelope of time-compressed speech in human auditory cortex (Nourski et al., 2009). Specifically, high-γ power, but not low-frequency, tracked the envelope of speech even when it was not comprehensible. As indicated in previous studies, high-γ power is susceptible to the regularity of auditory events (Eliades et al., 2014; Yellamsetty and Bidelman, 2018), which may reflect the neural activity more related to the acoustic feature. In contrast, θ phase envelope tracking was modulated by speech intelligibility to a large extent (Ding et al., 2014; Hauswald et al., 2020; Peelle et al., 2013). Our finding that high-γ power envelope tracking preferred the noise-vocoded speech, while θ phase envelope tracking preferred the natural speech (Figure 2), supports that these two bands may reflect distinct speech-specific processing.

Our data further showed that compared with high-γ power, θ phase feature decoding yielded a larger proportion of significant contacts and better decoding accuracy (Figure 3), indicating θ phase may reflect the speech processing related to the intelligibility. Our findings align well with a recent study (Synigal et al., 2020), showing that the envelope of the attended stimuli could be better reconstructed using low-frequency (1-15 Hz) compared that using high-γ power. Besides, there exists a relationship between the functions at different frequencies (Canolty et al., 2006; Wang et al., 2008; Zhang et al., 2020). Combining high-γ power and low-frequency feature could better reconstruct the speech envelope (Akbari et al., 2019; Synigal et al., 2020) and optimize selective neuronal tracking of the envelope of the attended speaker (Zion Golumbic et al., 2013). Thus, high-γ power and θ phase envelope tracking may play different roles in speech-specific processing.

### Hemisphere lateralization

Speech processing in the current perspective is proposed to be bilateral but with differential sensitivity to specific spectro-temporal features of speech (Flinker et al., 2019; Giroud et al., 2020). Based on the asymmetric sampling in time (AST) hypothesis (Poeppel, 2003), high- and low-frequency neural oscillations may have different hemisphere lateralization. Specifically, high-frequency neural oscillations with a short temporal integration window are lef-themisphere dominant, while low-frequency neural oscillations with a long temporal integration window are right-hemisphere dominant (Morillon et al., 2012). Similarly, our results revealed left-hemispheric lateralization in high-γ power envelope tracking and the right-hemispheric dominance in θ phase decoding accuracy. Since our data were obtained directly from human auditory cortex using sEEG technology with a high temporal-spatial resolution, this study provided prevailing evidence to the AST theory.

The lateralization of speech processing at different frequencies may reflect a spectral processing hierarchy (Giroud et al., 2020). From this view, the fast high-γ power matches the evoked onset response of the human auditory cortex (Elhilali et al., 2004), which may reflect a general response to the low-level acoustic features in speech. In contrast, the slow θ phase may further select the specific linguistic properties of the speech. Combined, these two stages of processing may shed light on the neural dynamics in cortical auditory processing in the left and right hemispheres.

In sum, we revealed two stages of speech envelope tracking, which are differently modulated by speech intelligibility. The current findings suggest that our auditory system may use different pathways to process sounds with and without semantic information, which may be an important mechanism for human brain to resolve speech information under adverse auditory environments.

## MATERIALS AND METHODS

### Participants

Ten patients (6 males and 4 females) with the mean age of 30.9 years old (from 13 to 46 years old), who were undergoing neurosurgical treatment for refractory focal epilepsy in the Beijing Tiantan Hospital and the Sanbo Brain Hospital of Capital Medical University, participated in this study. They were implanted with intracranial stereo-electroencephalogram (sEEG) electrodes based on clinical evaluation for respective surgery. All patients were righ-thanded with a self-reported normal hearing and provided written informed consent to participate. All patients were right-handed native Mandarin Chinese speakers. Experimental procedures were approved by the Ethics Committee of the Beijing Tiantan Hospital and the Sanbo Brain Hospital of Capital Medical University.

### Acoustic stimuli and experimental procedures

The natural speech used in this study was a Chinese “nonsense” sentence recorded from a young female talker (Gao et al., 2017). The “nonsense” sentences are correct in syntactical level but semantically meaningless, which were usually used as stimuli for speech intelligibility measurement (Yang et al., 2007). In the current study, the English translation of the natural speech is *“This kind of Suzhou has remembered your butterfly”* (“这种苏州已经记住你的蝴蝶”).

The spectrally degraded speech stimulus was a reconstructed version of the natural speech using a custom-designed 2-band noise-vocoder in MATLAB (Shannon et al., 1995). In detail, we first filtered the natural speech and a speech-spectrum noise (same power spectrum as the natural speech) into two bands (band 1: 80 to 1236 Hz; band 2: 1236 to 8820 Hz; spaced in equal steps along the cochlear frequency map) (Greenwood, 1990). Secondly, we extracted the envelope and the temporal fine structure (TFS) (Wang and Li, 2017) from the filtered signals of both the natural speech and a speech-spectrum noise. Thirdly, the envelope of each filter output from the natural speech was then multiplied by the TFS of the corresponding filter output from the speech-spectrum noise (Smith et al., 2002), resulting in two natural speech-noise chimeras. Finally, we summed these two chimeras (with energy normalized) to produce a noise-vocoded speech (NV). According to behavioral evidence (Newman and Chatterjee, 2013), noise-vocoded speech with 2 band is unintelligible. The correlation coefficient between the natural speech and the noise-vocoded speech was 0.99 (Figure 1).

Both the natural and the noise-vocoded speech were calibrated to the level of 70 dB SPL using a Larson Davis Audiometer Calibration and Electroacoustic Testing System (AUDit and System 824). The duration of either stimulus was 3.275 s. The acoustic stimuli were transferred using Creative Sound Blaster (Creative SB X-Fi Surround 5.1 Pro, Creative Technology Ltd, Singapore) and presented to participants binaurally at a sampling rate of 22050 Hz with insert earphones (ER-3, Etymotic Research, Elk Grove Village, IL). To avoid the prior knowledge effect of the natural speech on the noise-vocoded speech, we always presented the noise-vocoded speech first. Both stimuli were repeated 30 times. The inter-stimulus interval (ISI) between the two stimulus presentations was random between 1800 to 2200 ms.

Patients were reclining on a bed in their private clinical suite during the experiment. No seizures were observed at least 4 hours before the experimental procedure in any patients. We asked the patients to stay awake and passively listen to the sounds without any behavioral task. sEEG data were recorded with a Nihon Kohden (Tokyo, Japan) clinical monitoring system using a sampling rate of 2000 Hz with the online bandpass filter of 0.08-600 Hz in Tiantan Hospital or using the Nicolet video-EEG monitoring system (Natus Neuro, USA) with the sampling rate of 512 Hz (bandpass filter of 0.05-200 Hz) in Sanbo Hospital.

### sEEG contact localization

Each sEEG electrode (Huake Hengsheng Medical Technology Co Ltd, Beijing, China) had 8-16 macro contacts (0.8-mm diameter, 2.0-mm length, 1.5-mm apart from each other). All subjects underwent the pre-implant structural MRI (T1-weighted) and the post-implant CT scans. The T1 scan was firstly used to reconstruct the three-dimensional brain surfaces using BrainSuite (2018a) (http://brainsuite.org/), and then aligned to the post-implant CT using a rigid transformation in BioImage (http://bioimagesuite.yale.edu). We then identified the coordinates of sEEG electrodes using the Brainstorm toolbox (Tadel et al, 2011; http://neuroimage.usc.edu/brainstorm/) in the MATLAB environment. The anatomical of each contact was identified using an individual atlas (Joshi et al., 2020) of each patient. Contacts localized in the Heshl’s gyrus (HG), superior temporal gyrus (STG) were selected. The sEEG electrodes have carved out the way to directly investigate the function of human auditory cortex in auditory perception (Fang and Hu, 2021). All the subject-wised coordinates were then transferred into MNI coordinates for group plotting.

### Electrophysiological data analyses

#### Preprocessing

Raw sEEG data from each contact were imported into MATLAB (R2018a, MathWorks) and visually inspected for the presence of artifacts and interictal epileptic spikes (Lu et al., 2021), which were excluded from the further analyses. Raw sEEG data were pre-processed using the EEGLAB toolbox (Brunner et al., 2013) in the MATLAB environment. First, we extracted broadband signal (0.5-200 Hz) and high-γ signal (60-140 Hz) from the recorded sEEG data. Next, broadband signals, were segmented into single-trial epochs from - 1.3 s to 4.5 s around the sound onset. Meanwhile, the power envelope of the bandpass filtered high-γ signals were extracted by Hilbert transform, segmented into single-trial epochs from - 0.5 s to 4.5 s around the sound onset, down-sampled to 100 Hz and z-scored to the baseline. A contact was selected for the following analyses, if it had the significantly larger response power against baseline for a period lasting at least 10 ms (Bonferroni correction, *p* < 0.05) for both the broadband signal and the high-γ signal.

#### High-γ power envelope tracking

The tracking strength between neural activity (high-γ power) and the speech envelope was quantified by a cross-correlation function using the Matlab function *crosscorr*. Considering that the response lags the stimulus, we calculated the correlation coefficient between the averaged high-γ power and the stimuls envelope (down-sampled to 100 Hz) were calculated for the different time lags (0 to 200 ms in step of 10 ms, the stimulus relative to the response). Confidence intervals were obtained by assuming high-γ power and the stimulus envelope uncorrelated, if the maximum correlation coefficient at the optimal lag time was larger than the upper value of the confidence interval, it was identified as statistically significant and kept for further comparisons. Otherwise, the correlation coefficient was replaced by the zero value.

#### θ phase envelope tracking

We quantified θ phase envelope tracking using cerebro-acoustic coherence (CACoh) (Harding et al., 2019; Peelle et al., 2013), a measure of how sEEG signal phases align with speech envelope phases. Specifically, single-trial broadband signals were transformed to the time-frequency domain using the complex Morlet wavelets method with a frequency resolution of 0.5 Hz. The same time-frequency transformation was applied to the speech envelope. The phase coherence between single-trial broadband signal and the speech envelope was calculated for each of 14 frequency bins between 3 and 10 Hz according to the following formula:

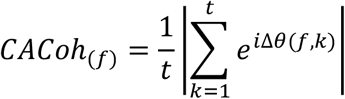

where Δ0(*f*, *k*) is the angular distance between speech envelope and sEEG signal at a single time point (*k*), *e* is the base of natural logarithms (~2.71), *i* is the imaginary unit that satisfies *i*^2^ = −1, and *t* is time.

Since cortical tracking of speech envelope lags the acoustic signal (Di Liberto et al., 2015; Ding and Simon, 2012), we first calculated CACoh for time lags up to 400 ms (10-ms steps) between the speech envelopes and single-trial sEEG signal. Then, for each contact, we obtained the peak CACoh value at the optimal lag for each frequency bin and averaged them over trials, resulting in one CACoh per frequency bin for the natural speech and noise-vocoded speech. To test the statistical significance of CACoh results, we we built a null distribution by randomly shifting the sEEG time series then calculating their CACoh 100 times. Actual CACoh values larger than 95% of the null distribution were considered significant, otherwise replaced by zero in further comparisons. We then calculated the z-score transform of CACoh values based on the null distribution and averaged the results in θ band (4-8 Hz).

#### Spectra-temporal decoding

To further investigate the relationship between the spectral contrast and neural oscillation activities, we used high-γ band power and θ-band phase to classify natural and noise-vocoded speech stimuli. We employed a support vector machine (SVM) approach using single trial time-frequency data from the high-γ power or θ phase, respectively. For high-γ power decoding, the responses (0 to 4000 ms, relative to the stimulus beginning) were segmented into non-overlapping 50-ms time bins, and these 50-ms windows were averaged to extract power values served as the model feature for this type. For θ phase decoding, responses were segmented into non-overlapping 100-ms time bins, and the model features were vectors of phase values by averaging across the 100-ms windows that were segmented. For each contact, we used the feature above separately to obtain model accuracy via a leave-one-out cross validation method. The SVM model was trained with a linear kernel and the penalty parameter set to 1. All analysis was performed in MATLAB (R2018b, MathWorks, MA, USA) using functions from the LIBSVM toolbox (https://www.csie.ntu.edu.tw/~cjlin/libsvm/). In order to test the significance of decoding values, we compared the observed decoding accuracy with a null distribution generated by permutation testing (swapping trial labels before SVM training). For each contact, the decoding accuracy with *p* value for high-γ band and θ band was obtained.

### Statistical analyses

Statistical analyses were performed with IBM SPSS Statistics 20 software (SPSS, Chicago, IL). Paired *t*-tests, independent-samples *t*-tests, chi-square test repeated-measures analysis of variance (ANOVAs) and post hoc comparisons (with Bonferroni corrections), and Pearson correlation tests were conducted. The null hypothesis rejection level was set at 0.05.

## Compliance and ethics

The author(s) declare that they have no conflict of interest.

## Acknowledgements

This study was supported by the National Key R&D Program of China (2017YFC1307500), National Natural Science Foundation of China (81771399, 32171039), Capital Healthy Development Research Funding (2016-1-2011, 2020-1-2013), and the Beijing Natural Science Foundation (Z200024).

